# A retrograde mechanism coordinates memory allocation across brain regions

**DOI:** 10.1101/2021.10.28.466361

**Authors:** Ayal Lavi, Megha Sehgal, Fardad Sisan, Anna Okabe, Donara Ter-Mkrtchyan, Alcino J Silva

## Abstract

Memories engage ensembles of neurons across different brain regions within a memory system. However, it is unclear whether the allocation of a memory to these ensembles is coordinated across brain regions. To address this question, we used CREB expression to bias memory allocation in one brain region, and rabies retrograde tracing to test memory allocation in connected presynaptic neurons in the other brain regions. We find that biasing allocation of CTA memory in the basolateral amygdala (BLA) also biases memory allocation in presynaptic neurons of the insular cortex (IC). By manipulating the allocation of CTA memory to specific neurons in both BLA and IC, we found that we increased their connectivity and enhanced CTA memory performance. These results – which are corroborated by mathematical simulations, and by studies with auditory fear conditioning – demonstrate that a retrograde mechanism coordinates the allocation of memories across different brain regions.

## Introduction

Memories are stored in sparse populations of neurons (i.e., memory ensembles or engrams) in multiple brain regions^1–3^. Previous studies have shown that memory ensembles in one brain region can affect ensembles in other brain regions involved in the same memory^4, 5^, and that neuronal oscillations may facilitate the coordination between these ensembles in different brain regions during memory encoding and retrieval^6–10^. A previous study also found increased structural connectivity between CA3 and CA1 memory ensembles in the hippocampus after memory formation^11^. Although there are mechanisms that regulate memory allocation to specific neurons in a given brain region^12–14^, it is unclear whether memory allocation is coordinated between the various brain regions involved in a memory.

Here, we report a novel approach to study the coordination between memory ensembles in different brain regions. We used this approach to study the allocation and coordination of memory ensembles across brain regions involved in conditioned taste aversion (CTA) and auditory fear conditioning (AFC). We find that biasing memory allocation in one brain region also biases memory allocation in connected neurons of other brain regions encoding the memory. In addition, we found that in task-relevant cortical and subcortical regions, memory ensembles in one region form direct monosynaptic connections with memory ensembles in other regions. Enhancing the connectivity between these memory ensembles resulted in an improvement in memory, a result that demonstrates the importance of these inter-regional memory networks. We also find that these changes in connectivity are specific to brain regions relevant to a given memory system (e.g., CTA vs. AFC). Our results, which we corroborated with mathematical simulations, demonstrate that in addition to anterograde mechanisms that propagate information across brain regions within a memory system^15–17^, retrograde mechanisms fine-tune the dynamic processes that coordinate the allocation of information and connectivity between memory ensembles in different brain regions.

## Results

### Tracing inter-regional memory networks

During CTA, temporally precise basolateral amygdala (BLA) activation is required to store taste memory traces in the insular cortex (IC) ^18–20^. We have previously demonstrated that viral expression of the cyclic AMP-responsive element-binding protein (vCREB) in a subset of neurons (vCREB neurons) in either the BLA or IC biases CTA memory allocation to those BLA or the IC vCREB neurons^14, 21^. We took advantage of CREB’s role in memory allocation, and rabies monosynaptic retrograde tracing^22, 23^ to develop an approach to study the connectivity between memory neurons across brain regions, a method that we call CRANE, for Cross-Regional Afferent Network of memory Ensembles. CRANE uses a combination of three viruses. This first virus carries the CREB gene which biases memory allocation to a sparse and random subset of neurons in a given brain region^12–14^ (Fig. 1a-b, AAV-RG-2A-CREB). The virus with CREB also has a rabies glycoprotein (RG) which is critical for rabies retrograde transsynaptic tracing ^22^. The second virus (AAV-TVA) includes TVA, a cell membrane protein that is essential for rabies infection^22^. Consequently, only cells with the RG-2A-CREB virus and with the TVA virus are capable of retrograde tracing with the rabies virus (Rabies-EnvA(PBG)ΔG-mCherry; Fig. 1a-b), the third virus in the CRANE system. Thus, the combination of these three viruses allows the tracing of monosynaptic connections to CREB-expressing neurons (i.e., likely to be memory ensemble neurons).

**Figure 1.**
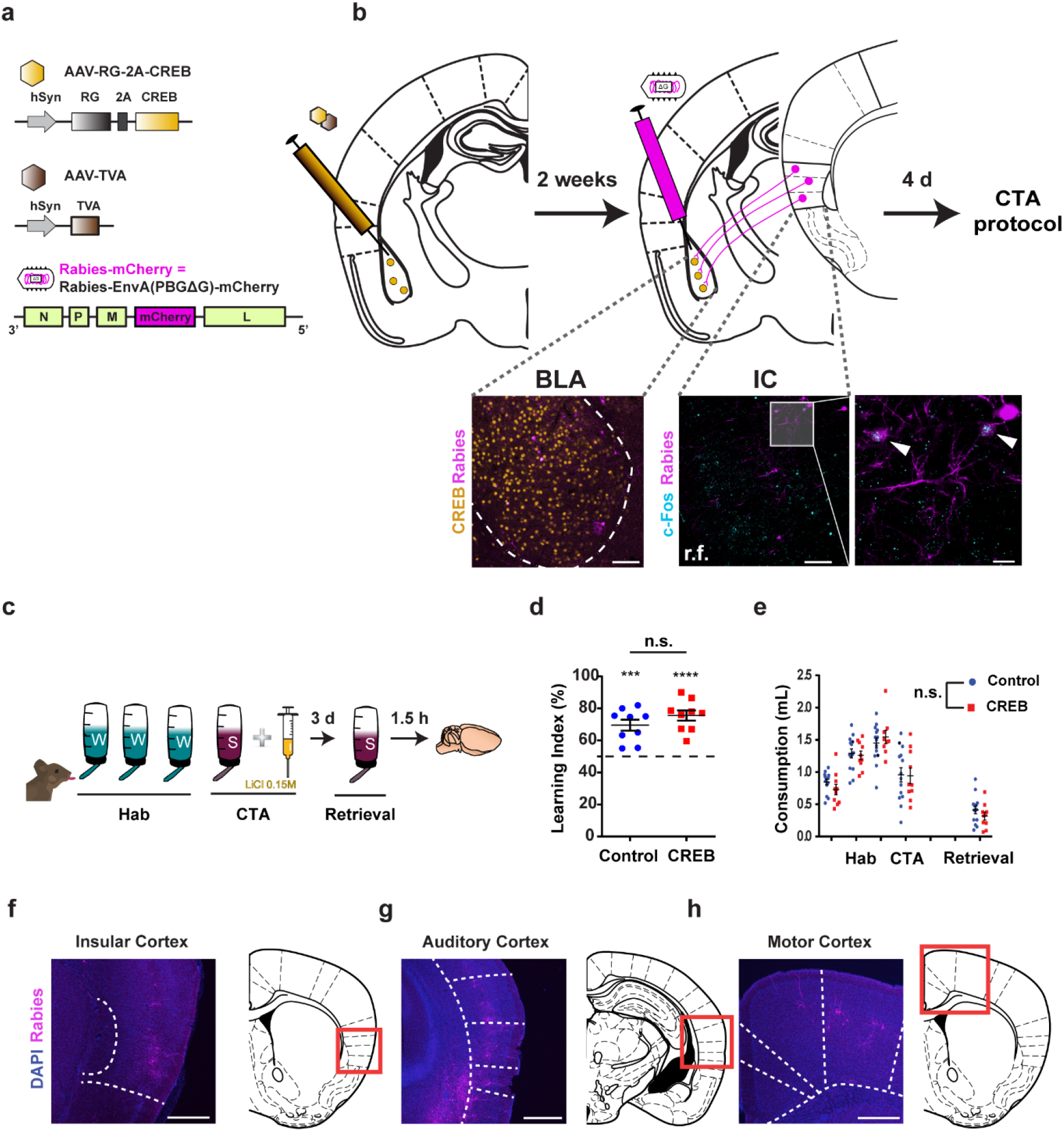
Using CRANE for tracing the connectivity of memory ensembles across brain regions. **a**, The CRANE strategy uses three viruses: an AAV virus, co-expressing CREB and the Rabies G protein (‘RG’) in tandem, biases the allocation of memory to specific neurons; RG together with TVA, from the AAV-TVA virus, targets a G-deleted Rabies-mCherry virus (‘Rabies-EnvA(PBG)ΔG-mCherry’) to the neurons with the AAV CREB virus (i.e., have a higher probability of becoming memory neurons). **b**, Two weeks after the injection of the two AAV viruses in the BLA, the G-deleted Rabies-mCherry virus was infused to the BLA. Following retrograde transport, and the CTA behavior protocol, rabies positive neurons (magenta, mCherry positive; connected to viral CREB expressing neurons in the BLA) expressing c-Fos (i.e., memory neurons) were identified in the IC, in the CREB and Control groups (white arrows; r.f. rhinal fissure). Scale bar 100 μm (20 μm for inset). **c,** Schematic diagram of CTA experimental design with 0.2% Saccharin (‘S’) and 0.15M LiCl (‘W’ = water). 1.5 hours following CTA retrieval, tissue was collected for immunostaining. **d,** Both RG-2A-CREB and the control RG-2A-CFP groups learn CTA (n = 9 mice per group, multiple one sample t-test, control \*\*\**P*<0.001 CREB \*\*\*\**P*<0.0001) with no difference between CREB and Control groups (unpaired t-test, t=1.293, *P* = 0.2143). **e,** Consumption of water during habituation (‘Hab’) and Saccharin during CTA acquisition (‘CTA’) and test (‘Retrieval’) was similar between both CREB and Control groups (2-way RM ANOVA, F(1, 22) = 0.2170, *P* = 0.6459). **f-h,** Representative images of trans-synaptically labeled neurons in several cortical brain regions. In all cases, the right panel is a stereotaxic map, with the red box showing the specific area for the image in the left panel. All of the regions with rabies positive neurons are known to project to the BLA. Scale bar 0.5 mm. All results shown as mean ± s.e.m.

The RG-2A-CREB and TVA viruses were injected into the BLA two weeks before the rabies virus was introduced. Since the rabies virus can only spread one synapse retrogradely^22^, and since it includes a mCherry gene, neurons that express mCherry throughout the brain (rabies positive neurons) should be monosynaptically connected to vCREB neurons in the BLA. We also determined whether these neurons, connected to vCREB neurons, are part of memory ensembles in other brain regions of the same memory system (i.e., whether they are part of an inter-regional memory network). In the experiments described here, one of the key controls we used included a virus where CREB was substituted by the cerulean fluorescent protein (CFP; AAV-RG-2A-CFP). As with the RG-2A-CREB virus, the RG-2A-CFP was randomly expressed in BLA. In this case, retrograde tracing allowed us to identify neurons connected to vCREB or control neurons in the BLA. Rabies virus preferentially traces active connections between neurons; therefore, rabies labelling has been shown to reflect experience-dependent changes in connectivity ^24, 25^. Consequently, our approach (CRANE) should preferentially trace learning-related connections between neurons in different brain regions.

Our CRANE studies in the BLA confirmed (Fig. 1f-h) well-known cortical projections to the BLA^26–30^, including those from the IC, the auditory cortex (AuCtx), and the primary motor cortex (M1). The brain regions identified by our experiments with the RG-2A-CREB and the RG-2A-CFP viruses were similar, showing that vCREB expression was not altering the brain regions connected to the BLA (Fig. S1b). Our neuroanatomic studies showed that RG-2A-CREB was expressed in a small sub-population (12.2 ± 2.78%) of the neurons in the BLA. As expected, following CTA training^14, 31, 32^, vCREB neurons in the BLA were 2 times more likely to be involved in CTA memory than both neighboring cells and chance levels (Fig. S1a, *P* < 0.01, t-test). This demonstrates that as expected, vCREB expression using our CRANE system was able to bias the allocation of memory to transfected BLA neurons. Thus, our CRANE system is well suited to track and study connectivity between memory ensembles across brain regions, including those activated by CTA, and involving the BLA.

### A retrograde mechanism coordinates memory allocation in inter-regional memory networks in the IC and BLA

To determine how memory ensembles in the BLA and the IC coordinate memory allocation during CTA, we used the CRANE approach in the BLA, and labeled memory networks with c-Fos immunostaining in the IC (Fig. 2a). Four days after rabies virus infusion, mice were habituated and trained on CTA (Fig. 1c), where a 0.2% Saccharin solution was paired with malaise-inducing LiCl. The group transfected with the RG-2A-CREB virus (CREB group) and the Control group (transfected with the RG-2A-CFP virus) both acquired CTA (Fig. 1d; *P* < 0.0001), showing a gradual increase in water consumption during habituation (Fig. 1e ‘Hab’), and a significant reduction in Saccharin consumption following conditioning (Fig. S2a ‘CTA’) and retrieval (Fig.S2a, two-way RM ANOVA, F(1, 21) = 51.35). Notably, no impairment in learning was detected following CREB or control virus injections (Fig. S2b).

**Figure 2.**
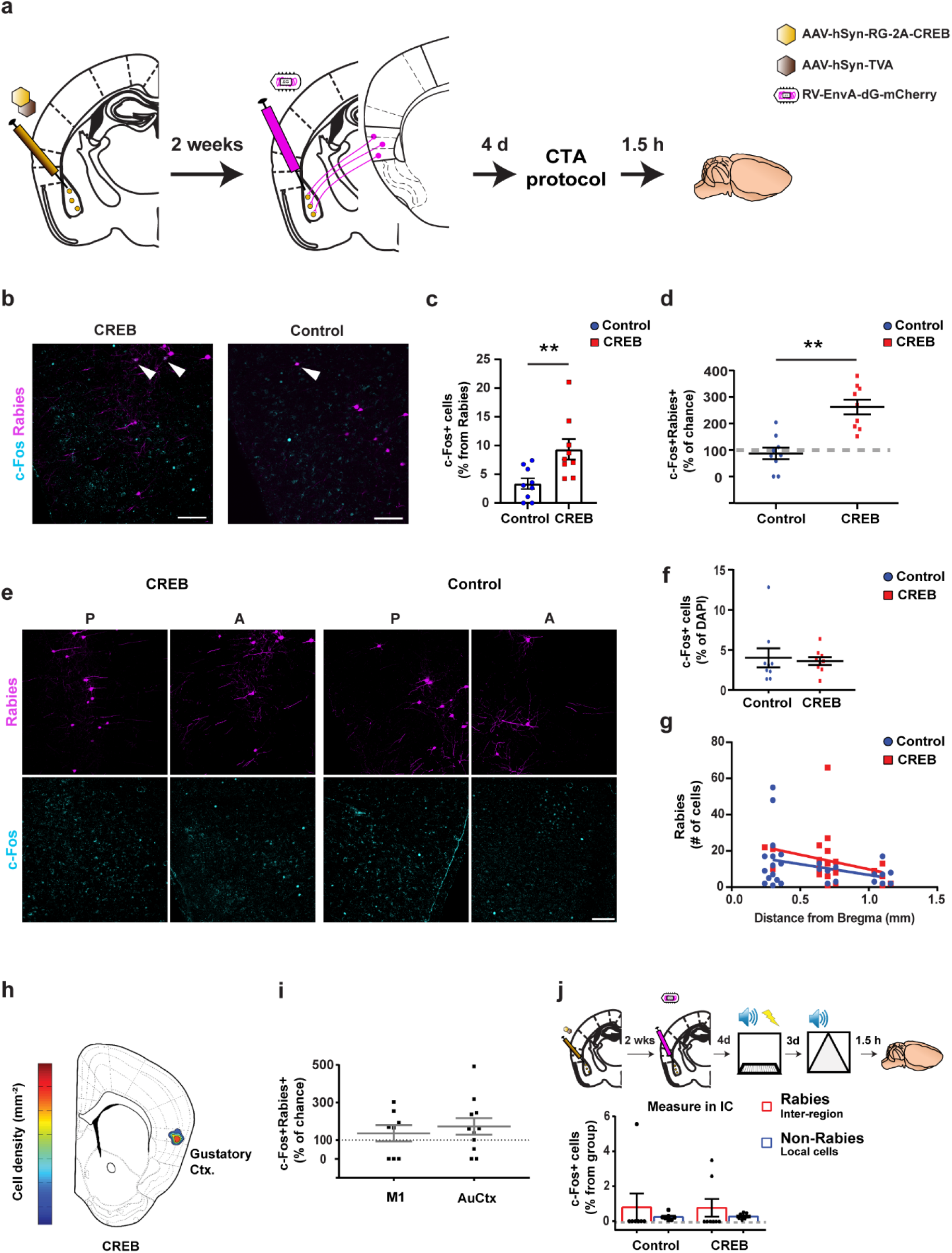
Connectivity between CTA memory ensembles in the IC and BLA. **a,** Schematics of the experimental design. RG-2A-CREB or the control RG-2A-CFP were injected in the BLA. **b,** Rabies-mCherry (rabies positive) and c-Fos expression mark presynaptic IC neurons that project to control or vCREB neurons in the BLA (arrows indicate co-expression of rabies and c-Fos; magenta and cyan respectively; scale bar 100μm). **c,** Presynaptic IC neurons innervating CREB neurons in the BLA are 2.5 times more likely to co-express c-Fos in comparison presynaptic IC neurons innervating CFP neurons in the BLA (n = 9 mice per group, unpaired t-test, **P=0.009). **d,** Analysis of the percentages of neurons in the IC that were both rabies positive and c-Fos positive for the CREB and CFP groups, normalized to chance levels per mouse. The CREB group, but not the CFP group (Control), showed a significant increase over chance levels of neurons in the IC that were both rabies positive and c-Fos positive (n = 9 mice per group, two-way ANOVA, F(1, 32) = 10.15, \*\**P*<0.01; Sidak’s multiple comparison test), showing that biasing memory allocation in the BLA, also biased memory in presynaptic neurons in the IC. **e,** Representative images of rabies and c-Fos expression along the Anterior-Posterior axis of the IC (‘A’ and ‘P’ respectively) in CREB and CFP (Control) group. Scale bar, 100μm. **f,** Similar expression levels of c-Fos in the IC in CREB and CFP (Control) mice following CTA retrieval. (n = 9 mice per group, unpaired t-test, P=0.758). **g,** The levels of rabies expression in the IC are similar along the A-P axis for CREB and CFP (Control) mice, showing that the efficiency of rabies tracing was comparable between these two groups (n = 27(7)/CREB 22(6)/control images(mice) ANCOVA test for equal slopes F(1,45)=0.372, P=0.545). **h,** Heatmap of the density of neurons in the IC that were both rabies positive and c-Fos positive (mm^−2^) in the CREB group; a hot spot (reflecting higher numbers of these neurons) is localized in the gustatory part of the insular cortex (‘Gustatory Ctx.’). **i,** Analyses of the percentage of neurons that were both rabies positive and c-Fos positive in CREB mice in brain regions not involved in CTA showed that this percentage was indistinguishable from chance levels, indicating that in these regions memory allocation was not biased to rabies positive neurons (M1 – primary motor cortex, AuCtx – auditory cortex; normalized to chance levels (n = 8-10 mice per group, two-way ANOVA F(1, 34) = 1.269, P=0.268). **j,** Task-specific expression of c-Fos in rabies positive IC neurons following auditory fear conditioning. No difference was found in c-Fos expression between the CREB group, the CFP (Control) group and chance levels (n = 7-8 mice per group, two-way ANOVA, F(1, 26) = 1.301, P = 0.2645). All results shown as mean ± s.e.m.

As mentioned above, our approach is designed to trace monosynaptic connectivity between memory ensembles across brain regions (Fig. 2b). Remarkably, our analyses showed that rabies positive neurons in the IC (expressing mCherry) in the CREB group were 2.5 times more likely to be activated (i.e., to be c-Fos positive) during CTA memory retrieval, when compared to the RG-2A-CFP virus control group (Fig. 2c, unpaired t-test *P* < 0.01), or to chance levels (Fig. 2d; two-way ANOVA F(1, 32) = 10.15, *P* < 0.01). Importantly, there was no difference between the groups in the overall percentage of c-Fos or rabies positive cells throughout the IC (Fig. 2e-g; the percent of rabies positive cells is in line with expected overall connectivity between IC and BLA^33–35^).

Notably, we found that changes in overall percentage of rabies positive cells in the IC reflected changes in connectivity following specifically CTA (i.e., above pre-CTA connectivity), since a similar CRANE analysis following auditory fear conditioning (AFC; i.e., does not involve the IC), revealed 71% reduction in the percentage of rabies positive cells in the IC (from 0.95±0.197% to 0.27±0.04%, Fig. S2c). These findings suggest that a retrograde mechanism has a role in tuning memory allocation and connectivity of projections from IC to BLA memory ensembles in CTA. This result is unexpected since previous studies assumed that anterograde mechanisms of information flow set memory networks throughout the brain^15–17, 36^.

Next, we carried out a number of experiments to test the behavioral and region specificity of the results just described. First, in addition to the IC, we also characterized rabies positive neurons and active neurons (i.e. expressing c-Fos) in primary motor cortex (M1), and in the AuCtx, two brain regions that are not involved in CTA^37, 38^. We found that the percentage of c-Fos positive neurons, amongst the rabies positive population in these two brain regions in the CREB group, was not different from the CFP control group, and from chance levels (Fig. 2i), a result that demonstrates that learning only engages presynaptic neurons in the brain regions relevant to the task (i.e., IC but not M1 or AuCtx for CTA).

Second, we also characterized rabies and c-Fos positive neurons in different subregions of the IC that are known to be involved in sensory modalities^39–41^ other than taste^33, 42^, including auditory processing^43^ and pain sensation^44^. Accordingly, we found that taste-specialized regions of the IC were the ones that showed the majority of the neurons that were both rabies and c-Fos positive (fig. 2h), a finding that attests to the brain region specificity of the c-Fos activation results described above.

Third, we repeated the CRANE BLA and IC experiment described above, except that we trained the mice in auditory fear conditioning (AFC). Unlike our results with CTA, our analyses showed that IC rabies positive neurons (mCherry positive) in the CREB group (BLA transfected with the RG-2A-CREB virus) were just as likely to be activated during AFC memory retrieval as the controls (BLA transfected with the RG-2A-CFP virus; Fig. 2j). Altogether these results demonstrate the behavioral and brain region specificity of the IC to BLA projections identified by CRANE, and reveals unprecedented evidence for the role of retrograde mechanisms in coordinating memory allocation and connectivity between the IC and the BLA memory ensembles.

### A retrograde mechanism coordinates memory allocation in bi-directional inter-regional memory networks in IC and BLA

Previous studies showed that the connectivity between the BLA and the IC is bi-directional^26, 27, 34–36, 41, 45^, and that projections in both directions mediate CTA encoding and retrieval^18, 34, 45, 46^. It is possible that the retrograde mechanism described above, that helps to coordinate memory allocation and connectivity from the IC to BLA during CTA, may also do so for BLA to IC connections. To test this hypothesis, we applied the CRANE approach described above to manipulate CTA memory allocation in the IC, and characterized c-Fos activation and rabies-related expression in the BLA (Fig. 3a). First, it is important to note that the CREB and the Control groups showed comparable levels of CTA (Fig. S3a; n = 8 mice per group, F(1,14)=42.42 for primary effect of retrieval, \*\*\*\**P* < 0.0001). As expected, vCREB neurons in the IC were 3 times more likely to express c-Fos in comparison to neighboring cells (Fig. S3d; T-test \*\**P* < 0.01), and to chance levels (\*\*\**P* < 0.001), confirming that CREB affects memory allocation in the IC^21^. We also confirmed that IC to BLA projections are more abundant than BLA to IC ones^26^ (Fig. S3c).

**Figure 3.**
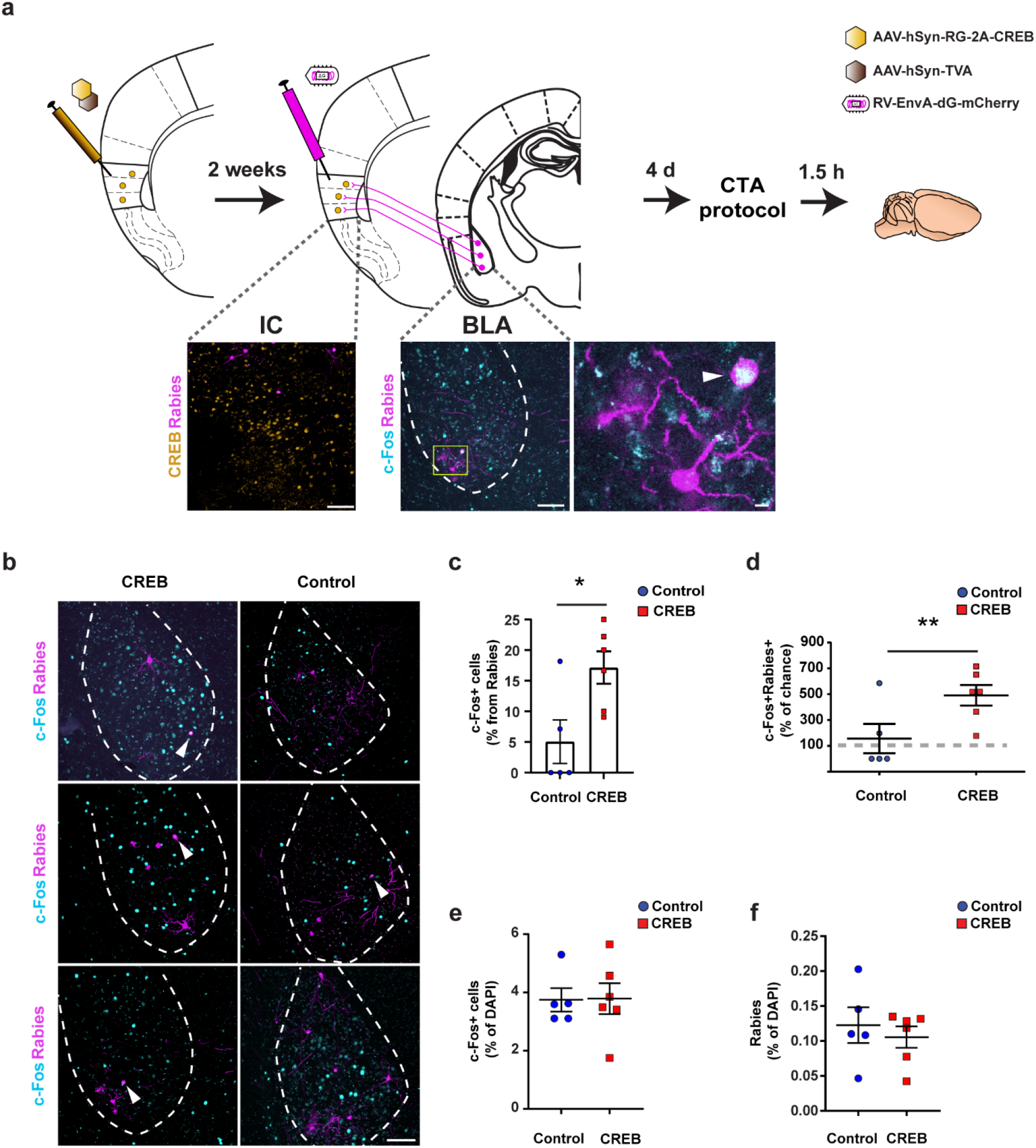
Bi-directional memory ensemble connectivity between the BLA and the IC. **a,** Schematics of experimental design. RG-2A-CREB or the control RG-2A-CFP were injected in the IC. Rabies-mCherry (rabies positive) and c-Fos expression mark presynaptic BLA neurons that project to control or vCREB neurons in the IC (yellow square in the left BLA image is magnified on the right, arrow indicates co-expression of rabies and c-Fos; magenta and cyan respectively). Scale bar 100 μm (20 μm for inset). **b,** Rabies-mCherry expression in the BLA co-localized with c-Fos immunostaining. CREB group shows consistently more co-localization between rabies and c-Fos positive cells in the BLA (white arrows). **c,** Presynaptic BLA neurons innervating vCREB neurons in the IC are 3 times more likely to co-express c-Fos in comparison presynaptic BLA neurons innervating CFP neurons in the IC (n = 5-6 mice per group, unpaired t-test, *P=0.021). **d.** Analysis of the percentages of BLA neurons that were both rabies positive and c-Fos positive for the CREB and CFP groups, normalized to chance levels per mouse. The CREB group, but not the CFP group (Control), showed a significant increase over chance levels of neurons that were both rabies positive and c-Fos positive in the BLA, demonstrating that biasing memory allocation in the IC biases memory allocation in presynaptic neurons of the BLA (normalization shows percent above chance level; two-way ANOVA, F(1,18)=7.597, *P=0.013; Sidak’s multiple comparison test \*\**P*<0.01). **e,f,** CREB and CFP control groups show same levels of c-Fos expression (**e**) and Rabies-mCherry (**f**) in the BLA following CTA (unpaired t-test, *P*>0.05). All results shown as mean ± s.e.m.

Importantly, our analyses showed that BLA rabies positive neurons (mCherry positive) in the CREB group were more likely to be activated (i.e., to be c-Fos positive) during CTA memory retrieval, when compared to the CFP virus control group (unpaired t-test, \**P* = 0.021; Fig. 3b,c), or to chance levels (Fig. 3d; 2-way ANOVA, F(1,18)= 7.597; \**P* < 0.05 for CREB vs. Control group, \*\**P* < 0.01 for CREB rabies vs. control rabies group). As expected, the percentages of overall rabies positive or c-Fos positive cells were not different between the CREB and Control group (Fig. 3e,f). These results show that the retrograde mechanism that we discovered coordinates memory allocation through projection from IC to BLA memory ensembles, also has a role in coordinating memory allocation in the projection from the BLA to the IC memory ensembles of this memory system. Together with the results presented above (Fig. 2), these findings demonstrate that there is a bi-directional monosynaptic coordination of memory allocation and connectivity in the IC and BLA^47, 48^. Additionally, these results also demonstrate the usefulness of CRANE in defining the mechanisms that coordinate memory ensembles across brain regions.

### A retrograde mechanism coordinates memory allocation in inter-regional memory networks in the AuCtx and the BLA

Beyond CTA, the BLA is involved in other forms of learning and memory^12, 14, 49^. For example, auditory fear conditioning (AFC) is known to engage the AuCtx and the BLA during memory encoding and retrieval^14, 49^. To test the hypothesis that a retrograde mechanism also plays a role in coordinating memory allocation between the BLA and AuCtx in AFC, we used the CRANE approach to manipulate AFC memory allocation in the BLA, traced connectivity, and characterized c-Fos activation in the AuCtx (Fig. 4a). Consistent with a role for CREB in AFC memory allocation in the BLA, we showed that during AFC memory retrieval, viral CREB-expressing neurons in the BLA were 3 times more likely to be activated (i.e., to be c-Fos positive) than control neurons (Fig. S4b, *P* < 0.01). Importantly, both the CREB and Control group showed evidence of AFC (Fig. 4b,S4a; no interaction, F_INTERACTION_(1,12) = 0.05 for the memory retrieval test).

**Figure 4.**
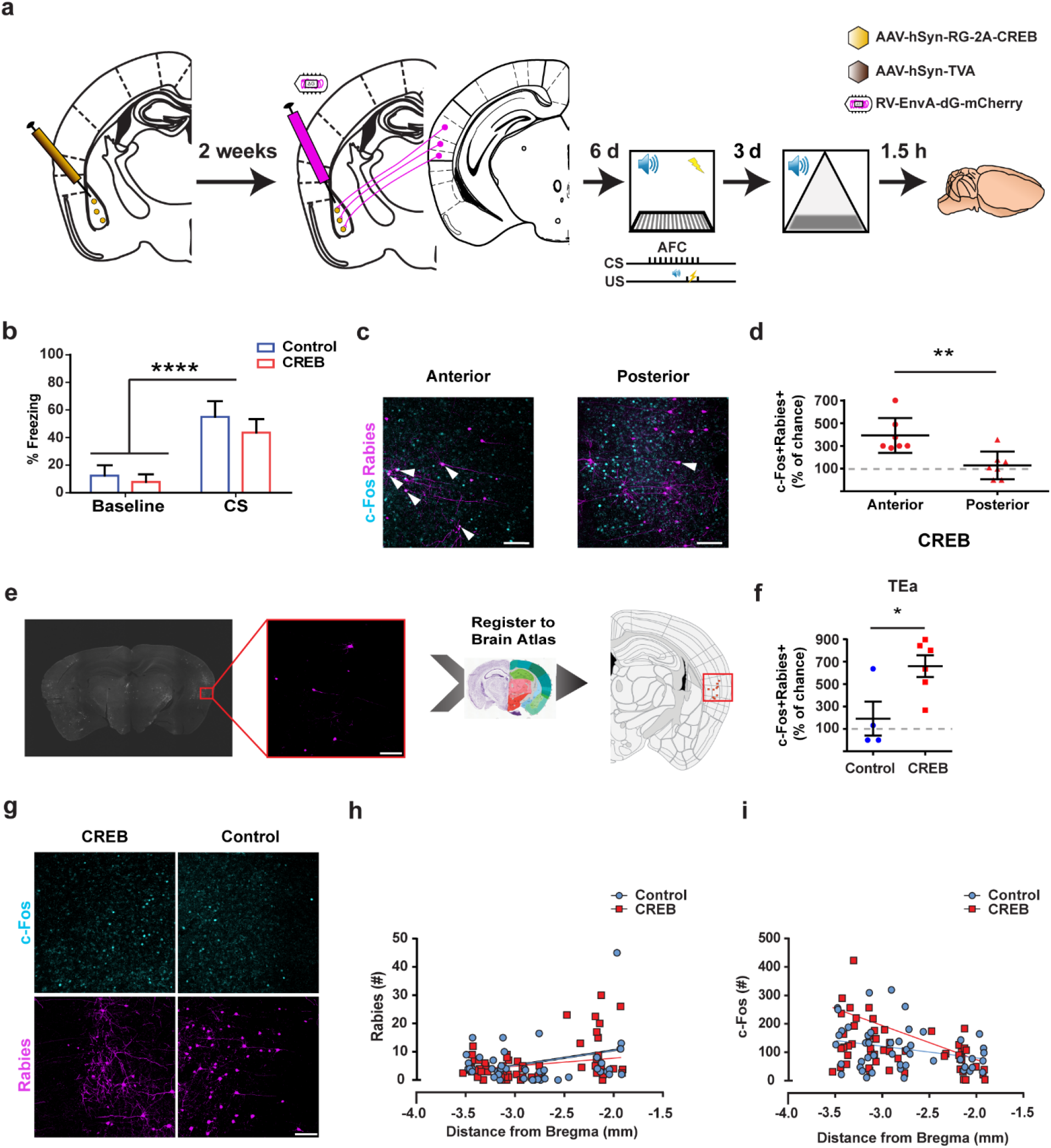
Inter-regional memory networks for auditory fear conditioning. **a,** AFC conditioning protocol comprises a series of 14kHz tones co-terminating with a foot-shock. This protocol has been previously shown to actively engage the auditory cortex ^53, 55^. **b,** Same levels of freezing following AFC protocol for CREB and Control group (two-way ANOVA, F(1,13) = 50.55, *P*<0.0001 for CS with no difference between groups). **c,** Representative images of activated rabies neurons (rabies positive and c-Fos positive neurons; white arrowheads) in anterior and posterior auditory cortex of the CREB group. The 14Hz tone has been previously shown to be encoded in the anterior auditory vortex ^53, 55^. **d,** In the CREB group, the anterior auditory cortex inter-regional memory network (rabies positive neurons) was highly activated in comparison to the posterior auditory cortex (n = 7 mice, unpaired t-test, \*\*\**P*<0.001) and to chance levels (multiple one sample t-test, CREB **P=0.001). **e,** Neuroanatomic-based analysis shows significant and specific activation (c-Fos+ Rabies+; representative image; scale bar 100 μm) in the temporal association area (**f,** ‘TEa’) of the auditory cortex (unpaired t-test, *P=0.0253; CREB also higher than chance level, multiple one sample t-test, **P=0.0022). **g,** Representative image from the auditory cortex in CREB and Control group. Scale bar 100 μm. **h,** Anterior auditory cortex shows slightly higher Rabies-mCherry expression in CREB and Control groups. Equal levels of Rabies-mCherry along the anterior posterior axis indicates the same level of rabies tracing for both groups. (ANCOVA test for equal slopes F(1,100)=0.972 P=0.3265). **i,** Equal levels of c-Fos expression between CREB and control group along the anterior posterior axis (ANCOVA test for equal slopes F(1,100)=0.923 P=0.339). All results shown as mean ± s.e.m.

The AuCtx displays a tonotopic map, where specific tones are preferentially activated in different parts of this structure^50, 51^. Previous studies^51, 52^ suggested that the specific tone used in our studies (14kHz) is encoded in the anterior AuCtx. Therefore, we analyzed the activation of rabies positive neurons (i.e., mCherry and c-Fos dual positive neurons) in the anterior vs. posterior AuCtx of the CREB group (Fig. 4c). We found that in the CREB group, rabies positive neurons in the anterior AuCtx, which is associated with the encoded tone^52, 53^, were more likely to be activated (i.e., higher percentage of c-Fos positive neurons) than both rabies positive neurons in the posterior AuCtx and chance levels (Fig. 4d, *P* < 0.001). In contrast, there were no differences among rabies positive neurons in anterior and posterior AuCtx (or chance levels) in the CFP control group (Fig. S4c).

Previous studies demonstrated that the connectivity and the tonotopic map changes along the dorsal to ventral axis of the AuCtx^27, 52^. Therefore, we registered the images we collected following AFC to the brain atlas^54^ (Fig. 4e,f) for an unbiased identification of regions with high levels of neurons that were both Rabies positive and c-Fos positive (c-Fos+ Rabies+). Analyses of the ventral area of the AuCtx (temporal association area), revealed that in this region rabies positive neurons in the CREB group were 7 times more likely to be activated (i.e., to be c-Fos positive) than either rabies positive neurons in the control CFP group (Fig. 4f, unpaired t-test, \**P* = 0.0253) or chance levels (multiple one-sample t-test, \*\**P* = 0.0022). This region is not only known to encode the specific tone we used for our AFC experiments, it has also been shown to be highly connected to the BLA^27, 43, 52, 55^. In contrast, in other regions of the AuCtx, we did not detect any differences between the groups and chance levels (Fig. 4g-i, S4d-e). These results demonstrate that the retrograde mechanism identified above not only has a role in coordinating memory allocation across brain regions in CTA, it also plays a role in coordinating memory allocation across brain regions for other types of learning and memory (i.e., AFC). As with CTA, we found that AFC cross-regional memory ensembles are highly anatomically and behaviorally specific. Altogether, these results suggest that retrograde mechanisms have a universal role in coordinating memory allocation across brain regions.

### Higher connectivity between memory networks is associated with enhanced CTA performance

Thus far, our findings indicate that in addition to allocation mechanisms that act locally within a brain region^12, 14, 21^, there are retrograde mechanisms that operate across brain regions to influence recruitment of presynaptic neurons into memory ensembles (see Fig. 2 and 3). To understand how these two processes work together to influence memory allocation within a circuit, we expressed vCREB in both BLA and IC simultaneously. Expression of vCREB in the BLA was part of the CRANE approach in order to assess the recruitment of BLA-projecting IC neurons (Fig. 5a). We reasoned that this creates two population of neurons within the IC that can compete for recruitment into the memory ensembles: vCREB neurons in the IC (where local allocation mechanisms operate) and IC neurons that project to BLA vCREB neurons (where retrograde mechanism operates). We asked if one of these mechanisms has a dominant role in recruiting IC neurons into the CTA memory ensemble.

**Figure 5.**
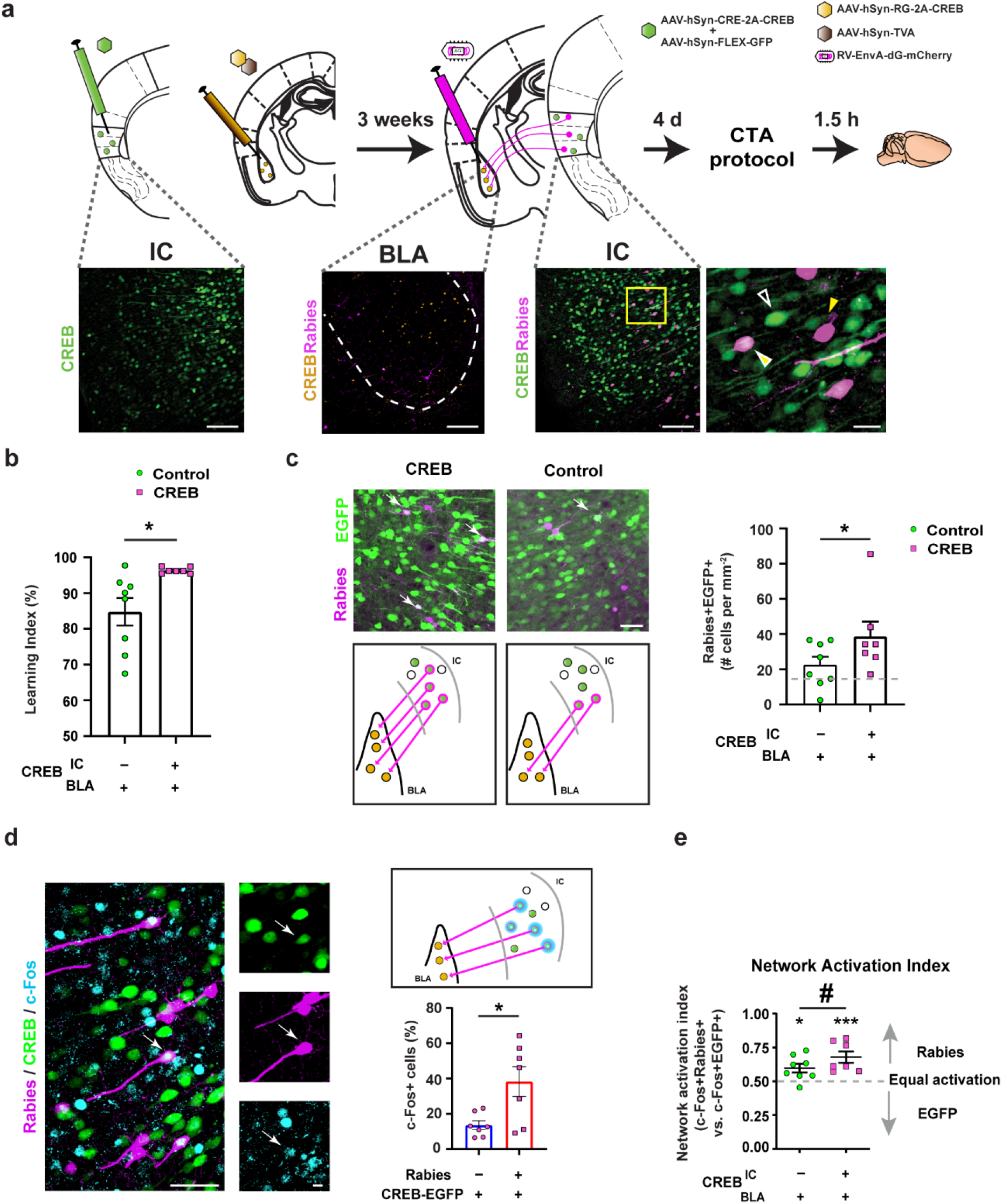
Simultaneous expression of viral CREB in the BLA and IC improves CTA performance by increasing the number and recruitment of BLA-projecting IC neurons. **a,** CREB virus was injected both in the BLA (orange, co-expressing RG) and the IC (green, co-expressing GFP, CREB^+^). Representative images from the IC show neurons expressing Rabies-mCherry (yellow arrow), CREB-GFP (hollow white arrow) and the combination of both (yellow arrow with white border). Three weeks later, Rabies was infused in the BLA. Scale bar, 100 μm (20 μm for inset). **b,** Expression of vCREB in BLA and IC enhanced learning to ceiling levels in comparison to the Control group (with vCREB expression only in BLA; L.I. 96 ± 0.3, control n = 8, CREB n = 7, unpaired t-test \**P*<0.05). **c,** Simultaneous vCREB expression within BLA and IC resulted in more projections from IC to BLA vCREB neurons. Top: Following CTA, we compared BLA projecting IC neurons (i.e., rabies-positive; ‘Rabies’) in the two groups (with and without vCREB expression within the IC). Neurons expressing both Rabies and GFP marked by white arrows). Middle: illustration demonstrating that in the CREB group, more IC vCREB neurons (green) are also rabies-positive (magenta borders) projecting the BLA vCREB neurons (orange), in comparison to the Control group (where GFP in expressed in the IC; green circles). Bottom: More vCREB neurons in the IC are rabies positive (GFP-mCherry positive - CREB group) in comparison to the control neurons (GFP-mCherry positive - Control group) indicating that viral CREB expression results in more BLA projecting vCREB neurons in the IC. These effects are also present when comparisons are made to chance levels (one-way ANOVA F(2,27)=5.148, *P=0.0128). **d,** Within the CREB group, vCREB IC neurons that project to vCREB BLA neurons are more likely to be recruited into the memory ensemble over their IC vCREB neighbors that do not project to the BLA. Top: Representative image of IC neurons expressing Rabies, CREB, and c-Fos (marked by the white arrow). Scale bar, 100 μm (20 μm for inset). Middle: Illustration demonstrating more activation (cyan circle) of IC vCREB neurons (green) that are rabies-positive (magenta borders) and project to vCREB neurons in the BLA, in comparison to neighbor IC vCREB neurons that do not project to the BLA. Bottom: During retrieval of the CTA memory, IC vCREB neurons that project to BLA vCREB neurons are more likely to be activated (GFP + mCherry + cFos positive neurons: CREB group) than IC vCREB neurons that do not project to the BLA (GFP + cFos positive neurons: CREB group). Unpaired t-test, *P*=0.0152. **e,** Comparison of c-Fos expression between CREB and Control groups: c-Fos expression in vCREB IC neurons (GFP + mCherry + cFos positive neurons: CREB group) or control IC neurons (GFP + mCherry + cFos positive neurons: control group) projecting to the BLA measured by network activation index (relative expression of c-Fos in Rabies (mCherry) neurons compared to GFP neurons per mouse; 0.5 = equal c-Fos expression levels in rabies positive and GFP positive cells). Both groups show that rabies positive neurons in the IC are more activated than local GFP/CREB positive neurons in the IC (one-way ANOVA F(2,20)=9.269, \*\*\**P*<0.001, \**P*< 0.05, #P<0.07). Scale bar, 100 μm (20 μm for inset). All results shown as mean ± s.e.m.

We found that vCREB expression both in the IC and the BLA resulted in enhanced CTA (Fig. 5b; t-test *P* = 0.02, *P* < 0.05), compared to vCREB expression only in the BLA (while maintaining vCREB function in biasing memory allocation; Fig. S5g). Next, we asked if this enhanced CTA was driven by local allocation mechanisms in the IC (recruitment of IC vCREB neurons) or the retrograde mechanisms (recruitment of IC neurons that project to BLA vCREB neurons). There were no differences in the overall levels of rabies, c-Fos, or GFP-positive neurons between the CREB and the Control group (S5b-d). Interestingly, we found that simultaneous vCREB expression within BLA as well as IC increased the connectivity between vCREB neurons across these brain regions. There was an increase in the number of vCREB neurons in the IC that are also rabies positive (GFP-mCherry positive - CREB group) in comparison to the control neurons (GFP-mCherry positive - Control group) indicating that expression of vCREB resulted in stronger connectivity between vCREB neurons (one way ANOVA F(2,27)=5.148 *P* = 0.0128; Fig. 5c and S5e). This level of connectivity was also more than expected by chance (Fig. S5f, t-test, *P* < 0.05). Hence, expression of viral CREB in the two brain-regions reorganized the vCREB ensembles such that they were more strongly connected.

We asked if this enhanced connectivity underlies better memory performance, specifically in the CREB group (seen in Fig. 5b and Fig. S5a; two-way RM ANOVA, F_CREB_(1,13) = 8.701 *P*=0.011). We found that within the CREB group, where vCREB is expressed in both IC and BLA, vCREB neurons in the IC that project to vCREB neurons in the BLA (GFP and mCherry-positive neurons) are more likely to be recruited into the memory ensemble (i.e, are cFos-positive) over their neighbors of vCREB neurons in the IC that do not project to the BLA (GFP positive and mCherry negative neurons; Fig. 5d, t-test *P* = 0.0152, *P* < 0.05). These data indicate that simultaneous expression of viral CREB in BLA and IC not only enhances the connectivity between vCREB neurons in these brain regions, but also increases the probability that these neurons will be recruited into the CTA memory ensemble. Since recruitment of IC vCREB neurons that project to the BLA (involving a retrograde mechanism) is more likely than IC vCREB neurons that do not project to the BLA (involving a local allocation mechanisms), we deduce that retrograde mechanisms play a dominant role in memory allocation within the IC. Consistent with our findings here, recent studies have shown that IC engram formation depends on BLA activity, thus suggesting that BLA memory ensembles help to shape engram formation in the IC^18, 20^.

In support of our hypothesis above, we found that IC neurons that projected to the BLA (rabies positive), whether or not they express viral CREB, were more likely to be activated (express c-Fos) than neighboring cells, (Fig. S5h, CREB \*\*\**P* < 0.01; Control \*\**P* < 0.001; two-way ANOVA F(1,26)=29.38, \*\*\**P* < 0.001). We also compared IC neurons that project to vCREB neurons in the BLA, but do not express vCREB themselves (Rabies-positive but CREB-negative), to IC vCREB neurons that do not project to BLA vCREB neurons (Rabies-negative CREB-positive). We found that the probability of activation (c-Fos expression) of the first group (Rabies-positive CREB-negative) was higher than the second group (Rabies-negative CREB-positive), both in the CREB group and in the control group (figure 5e, one way ANOVA F(2,20)=9.269 \**P* = 0.049, \*\**P* = 0.0011, #*P* = 0.0658), a finding that stresses the importance of retrograde mechanisms in memory allocation.

Altogether, these results show that increasing the connectivity between vCREB neurons in the IC and the BLA, also enhanced memory retrieval (Fig. 5b,c). We also presented multiple lines of evidence (Fig. 5d,e and Fig. S5e-g) that strongly suggest that the activation of memory ensembles in the IC is regulated by CREB levels within the BLA, providing additional lines of evidence for the role of retrograde mechanisms in coordinating the allocation and connectivity of memory ensembles across brain regions.

### A mathematical simulation supports the role of a retrograde mechanism in coordinating memory allocation across brain regions

The results presented above suggest that a retrograde mechanism has a key role in coordinating the allocation of memory between ensembles in different brain regions. To formally address this possibility, we ran mathematical simulations of three different possibilities that could conceivably account for the formation of memory ensembles in two interacting brain regions (i.e., BLA and IC; Fig. 6). This included an anterograde model, a random model and a retrograde model (see below). Four key elements were considered in these models, including BLA vCREB neurons, BLA-IC connectivity, BLA memory neurons (c-Fos positive neurons in the BLA), and IC memory neurons (c-Fos positive neurons in the IC; Fig. 6a). In these models, we used not only the data obtained from our results, but also data from the literature (see Methods). Importantly, we confirmed that the numbers obtained from our results, such as the percentage of memory (i.e., cFos positive) neurons in the IC, or the percentage of vCREB neurons, are consistent with published studies^12, 14, 21, 56^.

**Figure 6.**
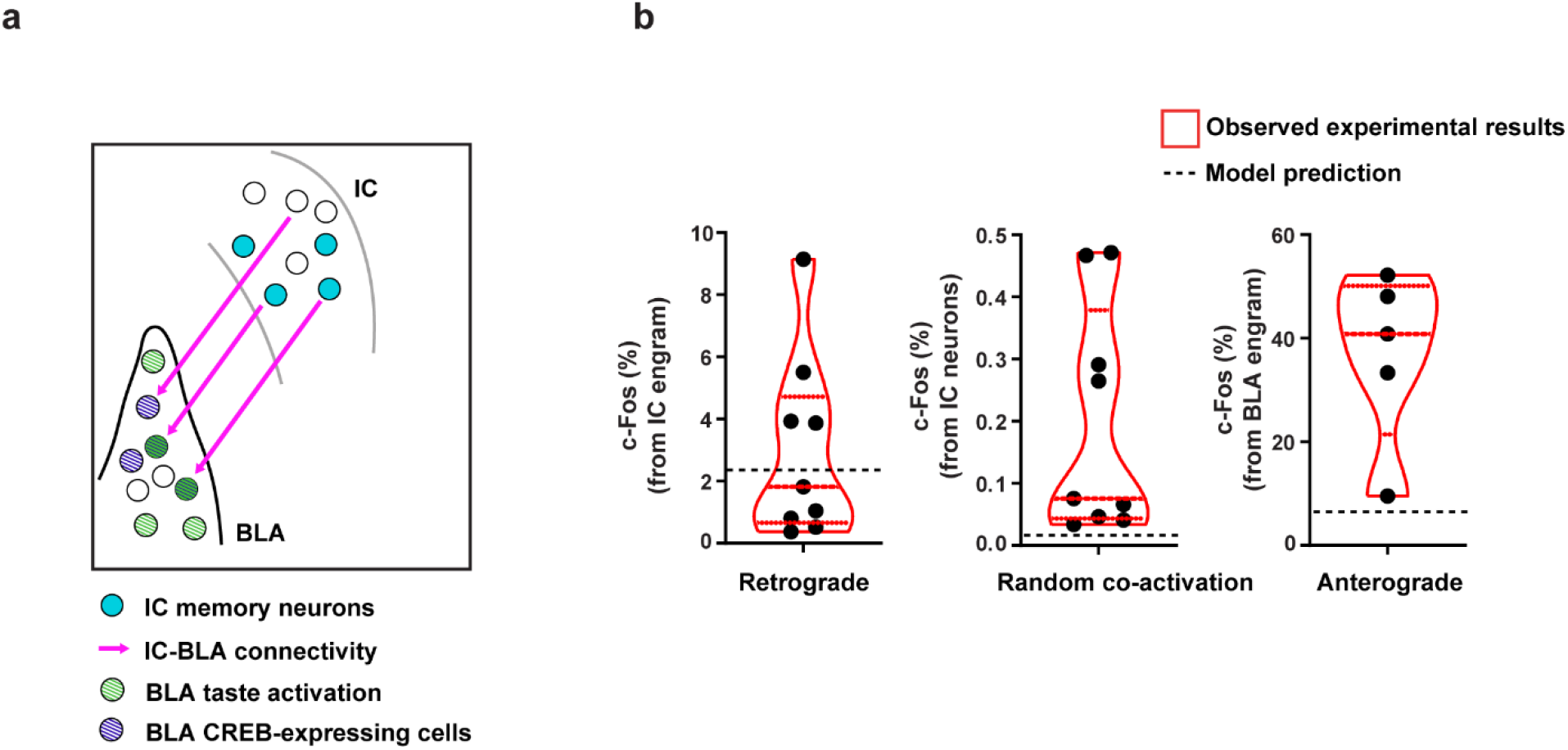
Retrograde model underlying cross-regional memory retrieval coordination. **a,** Illustration of three proposed models for cross-regional coordination of memory encoding and retrieval. The model considers the four key elements: BLA vCREB expressing cells, IC-BLA connectivity, BLA taste-specific cells, and IC memory neurons (using acceptable numbers from the literature, where available). **b,** Comparison of the predictions from the three models (black dashed line; model prediction) to experimental results (presented in violin plots. Dashed and dotted red lines within the violin indicate median, Q1, and Q3, respectively). (Retrograde) Comparing the expected activation of IC neurons projecting to BLA memory ensembles, we find the prediction falls well within the observed results (model prediction 2.22%, observed 3%). (Random co-activation) Predictions from the Random model, that assigns the cross-regional coordination to chance activation of connected neurons in both regions, show much lower expected activation in the IC than the observed activation (model prediction 0.029%, observed 0.2%). (Anterograde) The anterograde model predicts 6-fold-more activation of the BLA memory-encoding neuron in comparison to the observed activation (model prediction 36%, observed 6%).

The anterograde model is based on the idea that BLA neurons that are innervated by IC memory neurons have a higher chance of participating in memory encoding and storage. This idea, which has received tacit support in many neuroscience models^15–17^, assumes that information from the senses is processed in multiple downstream brain regions, and that memory neurons in one region connect to post-synaptic neurons in downstream regions, and thus determine which neurons in these downstream regions go on to encode and store memory. Based on our data and published studies ^14, 34, 57^, this model would predict (see Methods) that the percentage of vCREB positive BLA neurons that encode taste information (c-Fos positive), and receive presynaptic projections from IC memory neurons (Rabies positive and c-Fos positive), out of all the BLA memory neurons (c-Fos positive), should be 6% (Fig. 6b, right). In contrast, in our CRANE experiments, we found that 36% of the BLA memory neurons (c-Fos positive) were vCREB positive, a result that is 6 times higher than expected. This implies that an anterograde mechanism alone could not account for our results in the BLA.

In the random model, the allocation of neurons in the BLA and IC are independent of each other, and the connections between memory neurons across these two brain regions is thus determined by the random probability that taste-responding neurons in the IC would be connected to taste-responding neurons in the BLA. This model predicts that the percentage of IC memory neurons that project to BLA memory neurons should be 0.029% (Fig. 6b, middle). As with the anterograde model, we compared the predictions of this model to the observed percentage of IC memory neurons that connect to vCREB neurons in the BLA, and found that it is 0.2%. This is almost 7 times higher than what we observed, and therefore the random co-activation model alone could not account for our results in the BLA.

In the retrograde model, memory neurons in the BLA affect the probability that presynaptic neurons in the IC would be engaged in memory encoding and retrieval. This model predicts that the percentage of IC memory neurons that connect to vCREB BLA memory neurons, out of all of the IC memory neurons, would be 2.22% (Fig. 6b, left). Our observations confirm this prediction, since we found 3% ± 0.92% of IC memory neurons connect to vCREB BLA memory neurons, out of all of the IC memory neurons (Fig. 6b, left). These results show that the retrograde model best explains the results presented here. Altogether, our results demonstrate that the allocation of memory in different brain regions is coordinated and that a retrograde mechanism has a key role in this process. Consequently, these retrograde mechanisms also shape the connectivity of interregional memory networks, and this process affects memory formation and retrieval.

## Discussion

Here, we reported a novel approach (CRANE) to tag and trace the connectivity of memory networks across brain regions. Using CRANE and mathematical simulations, we showed that retrograde mechanisms help to coordinate memory allocation and connectivity across memory ensembles in different brain regions, and that this process is critical for memory. We found that IC neurons that are connected to BLA memory neurons, are more likely to be themselves involved in memory. These IC taste memory neurons are localized in the gustatory part of the IC (but not in other connected regions), suggesting that this retrograde mechanism engages specifically neurons in brain regions that are part of the same memory system. IC CTA memory neurons, that are connected to BLA memory neurons, are much more likely to be activated during memory recall than other IC neurons. This result highlights the role of the BLA in helping to shape memory networks in presynaptic IC neurons. We used vCREB in the BLA and IC to enhance the connectivity between these two brain regions, and showed that this could improve CTA memory. Importantly, we also found evidence that a retrograde mechanism also coordinates memory allocation in AFC ensembles in the BLA and AuCtx, suggesting that this retrograde mechanism is a general strategy for the coordination of memory allocation across different memory systems.

We provided a detailed characterization of the approach we introduced (CRANE) to tag and trace monosynaptic afferents of specific engram neurons. CRANE brings together a tool that guides memory allocation (by vCREB expression) with a tool for retrograde tracing (rabies virus). In agreement with previous studies^12, 14, 21^, we show that CREB biases memory allocation for both CTA and AFC in multiple brain regions. Additionally, rabies has been used for retrograde tracing across the brain, indicating that CRANE will be generally useful for tagging and tracing synaptic connections between memory neurons in different brain regions and uncover their inter-regional memory networks. Beyond tracing connectivity, rabies virus has also been used to track changes in plasticity induced by learning^24, 58^. Thus, retrograde tracing with rabies reflects both pre-existing connectivity and learning-induced changes in connectivity. Accordingly, our CRANE studies of CTA showed that 3 days after training there is a significant increase in the connectivity between the IC and the BLA, reflecting rapid plasticity-dependent changes in connectivity.

Memory systems involve multiple regions throughout the brain^1–3^. Previous studies showed that memory neurons within a brain region (hippocampus CA3 and CA1) can be directly connected^11^, and that memory neurons in one brain region can affect other memory neurons in a different brain region^4, 5^. Additionally, neuronal oscillations are thought to coordinate neuronal activity, facilitate communication between brain regions^6, 7^, and orchestrate brain-wide memory encoding and retrieval^8–10^. However, little is known about the principles (if any) that coordinate the allocation of memory between neuronal ensembles in these regions.

In taste processing, electrophysiological studies showed cooperative encoding of taste information in the BLA and the IC, which involves matching response dynamics, where the BLA displays earlier palatability-related information, and seems to drive responses in the IC^19, 59^. Additionally, BLA activity is required for CTA-mediated changes in the IC^18, 20^. As the IC and the BLA are bi-directionally connected, other studies further showed that BLA projecting neurons in the IC are modulated following CTA, and are critical for the formation and retrieval of aversive taste memory^33, 34^. The studies presented here show that a small subset of engram neurons in the IC and the BLA are monosynaptically connected, and are highly likely to be active during memory recall. This increased connectivity and enhanced activation during memory recall affected memory performance in CTA. These findings highlight the functional importance of this small set of interconnected neurons in the IC and the BLA. These findings also demonstrate the critical contribution of changes in connectivity to memory formation^4, 5^, and support previous studies demonstrating significant systems effects of small changes in neuronal networks^60, 61^.

Altogether the results reported here, as well as mathematical simulations, demonstrate that beyond anterograde mechanisms that set memory ensembles across brain regions, retrograde mechanisms have a role in shaping the allocation of memory in ensembles in different brain regions of the same memory system. We showed that these retrograde allocation mechanisms also shape the connectivity between memory ensembles in different brain regions. Importantly, we demonstrated the functional relevance of this retrograde mechanism, since we showed that enhancing the connectivity between memory ensembles in different brain regions (i.e., IC and BLA) resulted in an enhancement in memory, a result that also demonstrates the importance of coordinating these inter-regional memory networks.

## Materials and Methods

### Animals

Adult F1 hybrid (C57Bl/6NTac × 129S6/SvEvTac) mice 3 to 5 months old were used in behavioral analyses. Mice were group housed with free access to food and water, and maintained on a 12:12 hour light:dark cycle. All experiments were performed during the light phase of the cycle. All studies were approved by the Animal Research Committee at UCLA.

### Viral constructs

For the CRANE AAV system, Recombinant virus (rAAV5) for all plasmids, including pAAV-hSyn-CpBG-2A-HA-CREB (‘RG’) and pAAV-hSyn-TVA (‘TVA’) were prepared and purified as previously described ^62–64^. All AAV constructs were subcloned into pAAV-hSyn-hChR2(H134R)-EYFP; pAAV-hSyn-hChR2(H134R)-EYFP was a gift from Karl Deisseroth (Addgene plasmid # 26973; http://n2t.net/addgene:26973; RRID:Addgene_26973). For the Rabies helper viruses, we used CpBG as the Rabies G-protein helper. CpBG and TVA were a gift from Dr. Edward Callaway ^23^, and were sub-cloned together with the full-length CREB gene ^21^. For the pAAV-hSyn-iCRE-2A-HA-CREB, we subcloned iCRE from pDIRE into pAAV-hSyn-CpBG-2A-HA-CREB. pDIRE was a gift from Rolf Zeller (Addgene plasmid # 26745; http://n2t.net/addgene:26745; RRID:Addgene_26745). For control experiments, Cerulean fluorescent protein was subcloned from mCerulean-N1 into the respective constructs generating, pAAV-hSyn-CpBG-2A-HA-Cerulean and pAAV-hSyn-CpBG-2A-HA-Cerulean. Recombinant virus (rAAV5) was purified. Vector titers were determined by Real Time PCR. All titers for AAV viruses were above 1.6 × 10^12^ genome copies/ml.

### Surgery and virus infusion

Mice were anaesthetized with 1.5 to 2.0% isoflurane for surgical procedures and placed into a stereotactic frame (model #1900, David Kopf Instruments, Tujunga, CA) on a heating pad. Artificial tears were applied to the eyes to prevent drying. Subcutaneous saline injections were administered throughout each surgical procedure to prevent dehydration. In addition, carprofen (5 mg kg-1) and dexamethasone (0.2 mg kg-1) were administered both during surgery and for 2-7 days post-surgery. A midline incision was made down the scalp, and we used the stereotaxic drilling unit (model 1911, David Kopf Instruments, Tujunga, CA) to perform craniotomy. Water with amoxicillin was administered for two weeks. For virus injection, a Nanoliter injector (World Precision Instruments) was used to infuse virus with Micro4 Controller (World Precision Instruments). Virus was infused at 50 nL/min. After infusion, the capillary was kept at the injection site for 5 min and then withdrawn slowly. For rabies tracing, 300 nl of 3:7 volume mixture of pAAV-hSyn-CpBG-2A-HA-CREB (or pAAV-hSyn-CpBG-2A-HA-Cerulean as control) and pAAV-hSyn-TVA was injected into the BLA or IC. For the dual-CREB experiment, 300nl of 1:1 volume mixture of pAAV-hSyn-iCRE-2A-HA-CREB (or pAAV-hSyn-iCRE-2A-HA-Cerulean as control) and AAV8-FLEX-GFP was injected into the IC. Following virus injection for rabies tracing, stainless steel cannulas were implanted to improve recovery time from the virus infusion and reduce recurrent damage to the tissue. Bilateral guide cannulas (Plastics One, C313GS-5/SPC) were implanted and fixed on the skull with dental cement. After cannula implantation, mice were single-housed. Four days before the CTA protocol (6 for AFC), mice were mice were anesthetized and Rabies-EnvA(PBG)ΔG-mCherry (Salk Viral Vector core titer 1.0+E8; 0.8 μL, 100nL/min) was infused through the internal cannula (Plastics One, C313IS-5/Spc) at the helper viruses injection coordinates. After infusion, the internal cannula was left in place for an additional 8 min to ensure full diffusion. Coordinates used were taken from the mouse brain atlas ^65^ (relative to Bregma, midline, or dorsal brain surface and in mm): BLA: AP −1.3, ML ±3.3, DV −4.8; IC: AP +0.7, ML ±3.55, DV −2.8 and AP +0.2, ML ±3.7, DV −3.0.

### Conditioned taste aversion

The conditioned taste aversion task was carried out as previously described ^14, 31, 32^ with minor modifications. CTA training took place in the light part of the cycle, 2 weeks following surgery. Mice were water deprived for 24 h and then habituated to the training cage for 3 days to get their daily water ration within 30 min per day from two tubes (10 ml). Habituation started 4 days after rabies virus infusion. On the conditioning day, the two tubes were filled with 0.2% saccharin sodium salt (w/v, the taste CS) instead of water. The CS was presented for 30 min and 20 min later, mice were treated with the malaise inducing agent lithium chloride (LiCl; 0.15 M, 2% body weight i.p.). Testing for aversion to saccharin occurred 3 days later. Two tubes (containing saccharin) were presented for 30 min. The intake of each fluid was measured and the learning index (LI) was defined as follows: [milliliters consumed during training/(milliliters during training + milliliters during retrieval)]×100%. For the dual-CREB experiment, we used 0.1% saccharin to uncover changes in LI and avoid ceiling effects. For this experiment LI was calculated: [milliliters of water consumed/(milliliters of water + milliliters of saccharin consumed)]×100%. 50% LI is equal preference level, and the higher the LI, the less mice preferred saccharin). We confirmed that both LIs were correlated (Pearson correlation; R^2^ = 0.961, *P* < 0.0001).

### Auditory fear conditioning

Auditory fear conditioning was carried out as previously described^14^ with minor modifications. Training consisted of placing the mice in a conditioning chamber,and 2 min later presenting a series of 10 tones, 14kHz each at 85dB, co-terminating with a 2-sec electric foot shock (0.5mA) through the floor grid. This was repeated 5 times at pseudo-random intervals (30 to 60 sec), and then mice remained in the chamber for an additional 2 min. Test for auditory fear conditioning occurred 3 days later. Mice were placed in a novel chamber (Context B) and 2 min later the tone CS was presented (for 1 min). Our index of memory, freezing (the cessation of all movement except for respiration), was assessed via an automated scoring system (Med Associates Inc.) with a 30 frames/sec sampling; the mice needed to freeze continuously for at least 1 sec before freezing was counted.

### Tissue processing and Immunostaining

Mice were transcardially perfused 90 minutes following behavioral retrieval with 4% PFA (4% paraformaldehyde in 0.1 M phosphate buffer), and after perfusion brains were extracted and incubated with 4% PFA overnight at 4°C. Coronal sections were cut at 50 μm on a microtome and transferred to PBS, then blocked in 5% Normal Goat Serum in 0.1 M PBS and 0.1% TritonX-100 for 1 hr. After blocking, sections were incubated in a primary antibody mix (in 0.1 M PBS, 0.2% TritonX-100 and 5% Normal Goat Serum) of rabbit anti-c-Fos (Cell Signaling, #2250,1:700) for two days at 4°C. After 3 × 15 min washes in 0.1 M PBS and 0.2% TritonX-100 the secondary antibodies were applied (in 0.1 M PBS, 0.2% TritonX-100 and 3% Normal Goat Serum): Alexa647 goat anti-rabbit (Invitrogen #A-21245, 1:1000 dilution). Slices were incubated in the secondary mix for 2 hr at room temperature. After 2 × 15 min washes in 0.1 M PBS and 0.1% TritonX-100, slices were incubated with 4’,6-diaminodino-2-phenylindole (DAPI, Life Technologies D-21490, 1:2000) for 15 min, and then were further washed with 0.1 M PBS and 0.1% TritonX-100 for 15 min before mounted onto slides with ProLong Gold antifade mounting media (Life Technologies, P36934). All immunostaining images were acquired with a Nikon A1 Laser Scanning Confocal Microscope (LSCM). Whole-slice images were obtained by tiling 4x images. Analysis was performed on 20x Z-stack images.

For quantification of c-Fos and mCherry (for Rabies), we used the Nikon Elements 3D counting module. All counts and co-localizations of c-Fos and rabies were verified manually by an expert blind to image identity. For Fig. 2h, heatmap was generated base on localization of cells that were both rabies positive and c-Fos positive in the Allen Mouse Common Coordinate Framework brain atlas ^66^. To normalize for chance, we subtracted chance (rabies/DAPI) × (c-Fos/DAPI) × 100 from the observed overlap (rabies and c-Fos)/DAPI × 100 and then divided by chance. For Fig. S5e, calculation was normalized to the baseline of GFP positive and c-Fos positive, per mouse. To calculate the Network activation index (Fig. 5i), chance levels were calculated for rabies (*chanceRabies*) and GFP (*chanceGFP*) Then, the difference-sum ratio was calculated DR = (*chanceRabies - chanceGFP)/(chanceRabies – chanceGFP+2*) and scaled to an index between 0 and 1 NAI = (DR+1)/2.

### Whole-brain analysis

Brain slice images were aligned automatically to the corresponding Bregma, based on the Allen brain atlas, and then brain maps were fine-tuned manually by an expert based on anatomic landmarks in an unbiased manner. Next, previous cell count data that were obtained from Z-stack analyses were overlaid on the brain Atlas using the WholeBrain platform^67^ and a custom-made R script.

### Model construction and parameters

We propose 3 models that could account for how memories are formed between directly connected neurons across brain regions. All models assumed that each BLA neuron is innervated by at most one neuron from the IC. We used numbers from the literature, where available. These include the number and proportion of neurons in the BLA and IC^68–70^, the extent of projections from IC to BLA^34, 35^, c-Fos in the BLA following CTA retrieval^71^, and the extent of neuronal response to taste in the BLA^57, 59, 72, 73^ and the IC^33, 74–76^. We also confirmed that several experimental numbers we used match measurements from previous studies, such as the percentage of neurons in the IC that are activated upon CTA retrieval^21^ and the percentage of CREB-expressing neurons in the BLA^12, 14, 56^.

#### The Anterograde Model

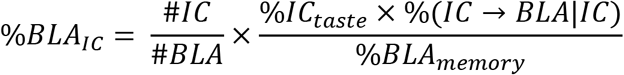

The anterograde model simulated the percentage of neurons in the BLA that encode taste and received projections from GC memory neurons out of all BLA memory neurons (%BLA_IC_). Activation of neurons in the IC, that are directly connected to the BLA, induces the activation of their counterparts in the BLA. Taking into account the ratio between the number of neurons in the IC (#IC) and the number of neurons in the BLA (#BLA), the anterograde model assumes that only BLA neurons that are directly connected by IC neurons participate in memory encoding and its subsequent storage. This model takes into account the percentage of taste-activated neurons in the IC (%IC_taste_) and the connectivity from the IC to the BLA (%IC→BLA|IC), out of the memory neuronal population in the BLA (%BLA_memory_). The model predicts that the percentage of BLA neurons that encode taste and receive projections from IC memory neurons, out of all the BLA memory neurons, is 6%. Since in our experimental setup, CREB neurons are the ones connected to the IC (and CREB is expressed in random cells prior to the CTA), we compared the prediction of this model to the observed percentage of BLA CREB-expressing neurons that were activated during learning. We found that 36% of the BLA memory neurons were CREB – 6 times higher than expected. This implies that the anterograde mechanism in of itself would not be sufficient to explain our results in the BLA. (for this model we used the following numbers: #IC = 90,000, #BLA = 194,000, %IC_taste_ = 36%, %IC→BLA|IC = 0.5%, %BLA_memory_ = 1.38%)

#### The Random Model

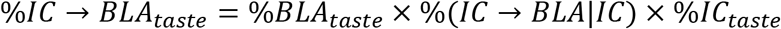

The random co-activation model simulates the percentage of IC neurons that encode taste and project to BLA neurons that are also activated by taste (%IC→BLA_taste_). According to this model, memory allocation in both the BLA and the IC is random and the connection between memory neurons across these brain regions is thus determined by the percentage of taste-responding neurons in the IC (%IC_taste_) that are connected (%IC→BLA|IC) to taste-responding neurons in the BLA (%BLA_taste_). This model predicts that the percentage of IC neurons that encode taste and project to BLA neurons, that are also activated by taste, is 0.029%. As with the Anterograde model, we compared the predictions of this model to the observed percentage of IC neurons that connect to CREB-expressing neurons in the BLA and found that it is 0.2% (almost 7 times higher than what we observed). (for this model we used the following numbers: %IC_taste_ = 36%, %IC→BLA |IC = 0.5%, %BLA_taste_ = 16%)

#### The Retrograde model

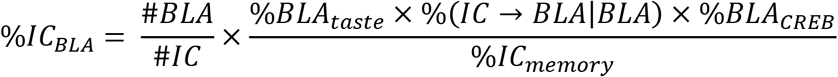

The Retrograde model simulates the percentage of IC neurons that encode taste and project to BLA memory neurons out of all IC memory neurons. In this model, memory neurons in the BLA affect the probability that IC neurons that project to them participate in memory encoding and retrieval. This model considers the probability that CREB-expressing neurons (%BLA_CREB_) that are activated by taste (%BLA_taste_) and receive projections from the IC (%IC→BLA|BLA) out of the memory neuronal population in the IC (%IC_memory_). The model predicts that the % of IC memory neurons that connect to CREB-expressing BLA memory neurons, out of all of the IC memory neurons, would be 2.22%. Our observations confirm this prediction since we found 3% ± 0.92% of such IC neurons, well within the expected experimental error. (for this model we used the following numbers: #IC = 90,000, #BLA = 194,000, %BLA_taste_ = 16%, %IC→BLA|BLA = 1.71%, %IC_memory_ = 3.6%, %BLA_CREB_ = 13.56%)

### Quantification and Statistical Analyses

The investigators who collected and analyzed the data, including behavior and staining, were blinded to treatment conditions. Error bars in the figures indicate the SEM. All statistical analyses were performed using GraphPad Prism 9. N designates the number of mouse or brains collected, unless otherwise stated. We analyzed at least 4 images per mouse from both left and right hemispheres. Statistical significance was assessed by Student’s t test, or one- or two-way ANOVA where appropriate, followed by the indicated post-hoc tests. The level of significance was set at P < 0.05.

## Acknowledgements

We thank C. Bear, A. Chien, D. Vuong, for advice and technical support. This work was supported by grants from the NIMH (R01 MH113071), NIA (R01 AG013622), and from the Dr. Miriam and Sheldon G. Adelson Medical Research Foundation to AJS.

## Author Information

The authors declare no competing financial interests. Correspondence should be addressed to A.J.S.

## Supplementary figures

**Figure S1.**
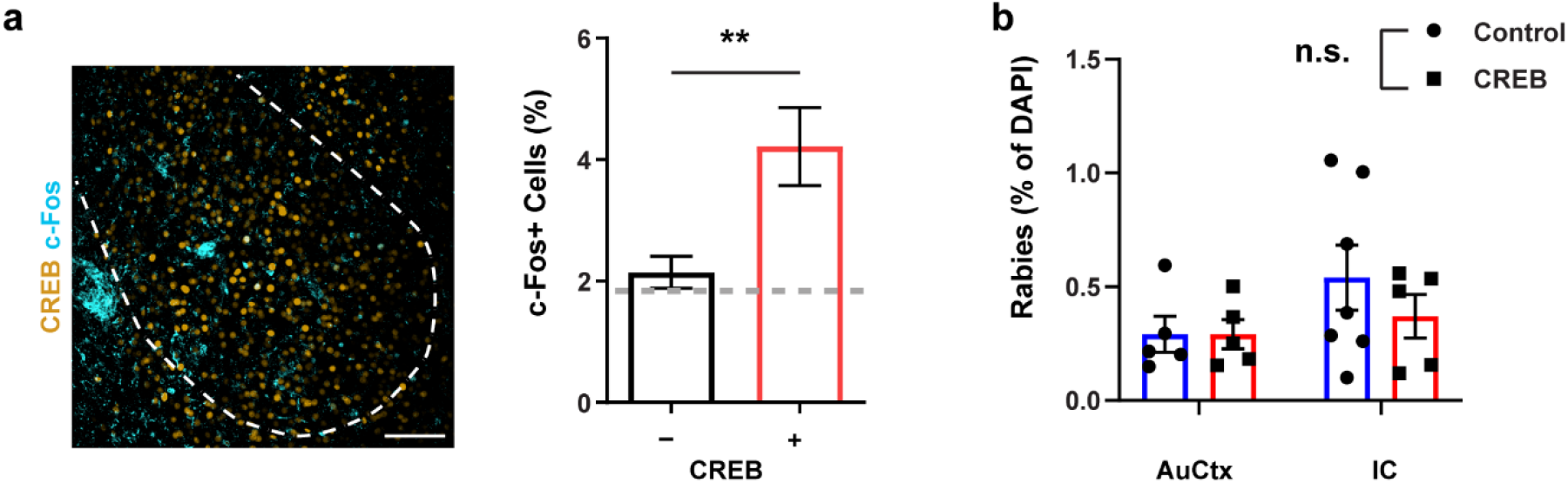
Characterization of the cross-regional trans-synaptic memory tracing expression system. **a,** Analysis of c-Fos expression in CREB neurons in the BLA (Representative image; cyan = c-Fos, orange = CREB). Comparison of c-Fos expression in CREB-expressing neurons (CREB +; 4.2%) and neighboring neurons (CREB -; 2.1%) shows bias in memory allocation to CREB neurons. (n = 13 images/5 mice, Pooled unpaired two tailed t-test \*\**P*<0.01). Dashed line indicates chance levels. Scale bar 100 μm. **b.** No difference between CREB and control (CFP) groups in the expression of Rabies (%Rabies of DAPI) in the auditory cortex (AuCtx) and the insular cortex (IC) (IC n = 7, AuCtx n = 5; two-way ANOVA F_Interaction_ (1, 18) = 0.5590. *P* = 0.464). All data in the figure are shown as mean ± s.e.m.

**Figure S2.**
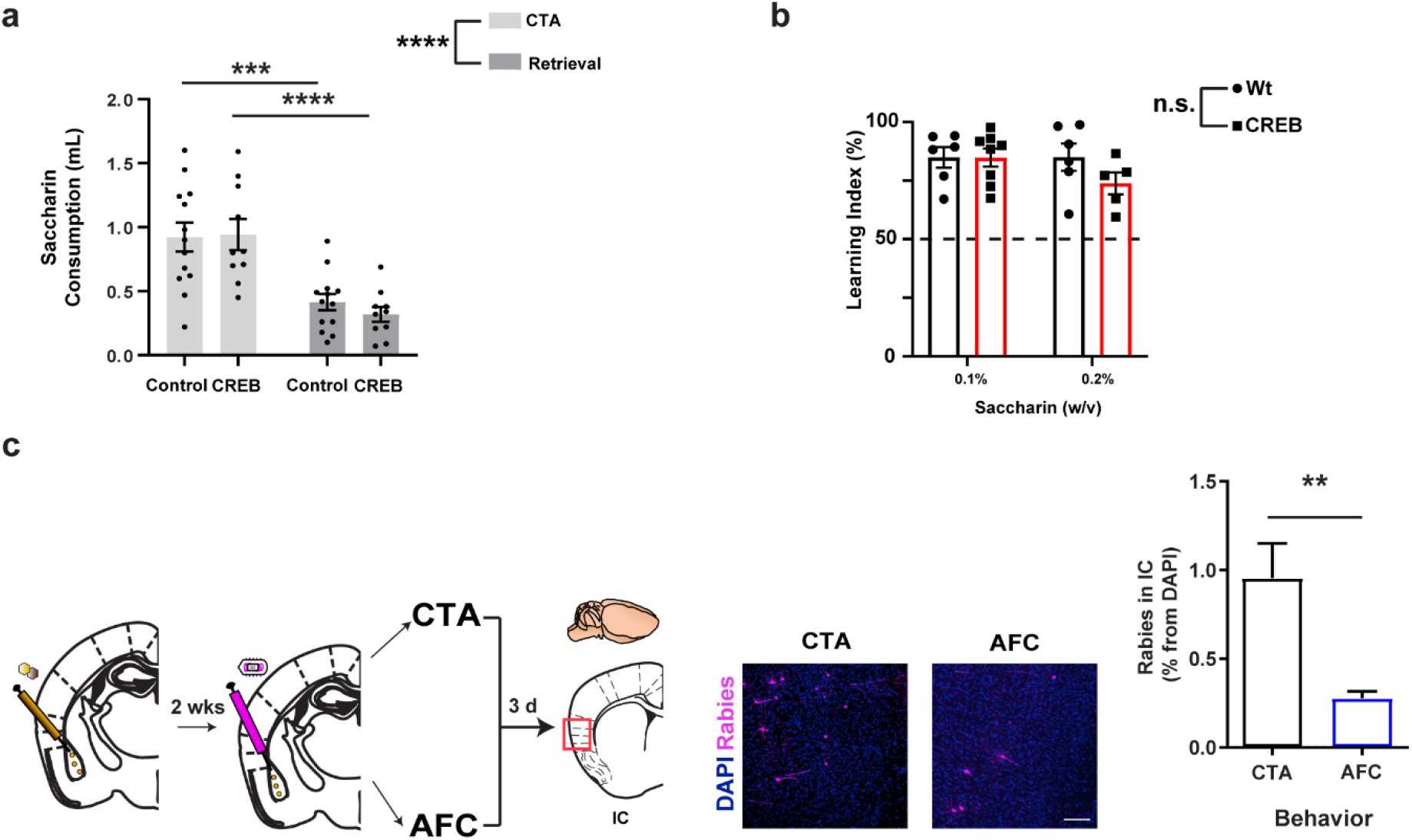
Characterization of CTA learning and retrieval with the CRANE system. **a**, Mice reduce Saccharin consumption following CTA in both CREB (n = 10) and control (CFP, n = 12) groups (two-way RM ANOVA, F(1, 21) = 51.35, \*\*\*\**P*< 0.0001, \*\*\**P*<0.001). **b**, CREB expression did not affect the learning index, regardless of Saccharin concentration (Two-way ANOVA, F(1, 18) = 2.072, *P* = 0.1672). **c,** Mice were trained in either CTA or auditory fear conditioning (AFC). Then, 3 days later they were given a retrieval test; Percentage of rabies positive neurons from DAPI staining measured in IC after either the memory retrieval test for CTA or AFC (CTA n = 25 AFC n = 15; unpaired t-test, **P = 0.0023 with Welch’s correction). Scale bar 100 μm. All results shown as mean ± s.e.m.

**Figure S3.**
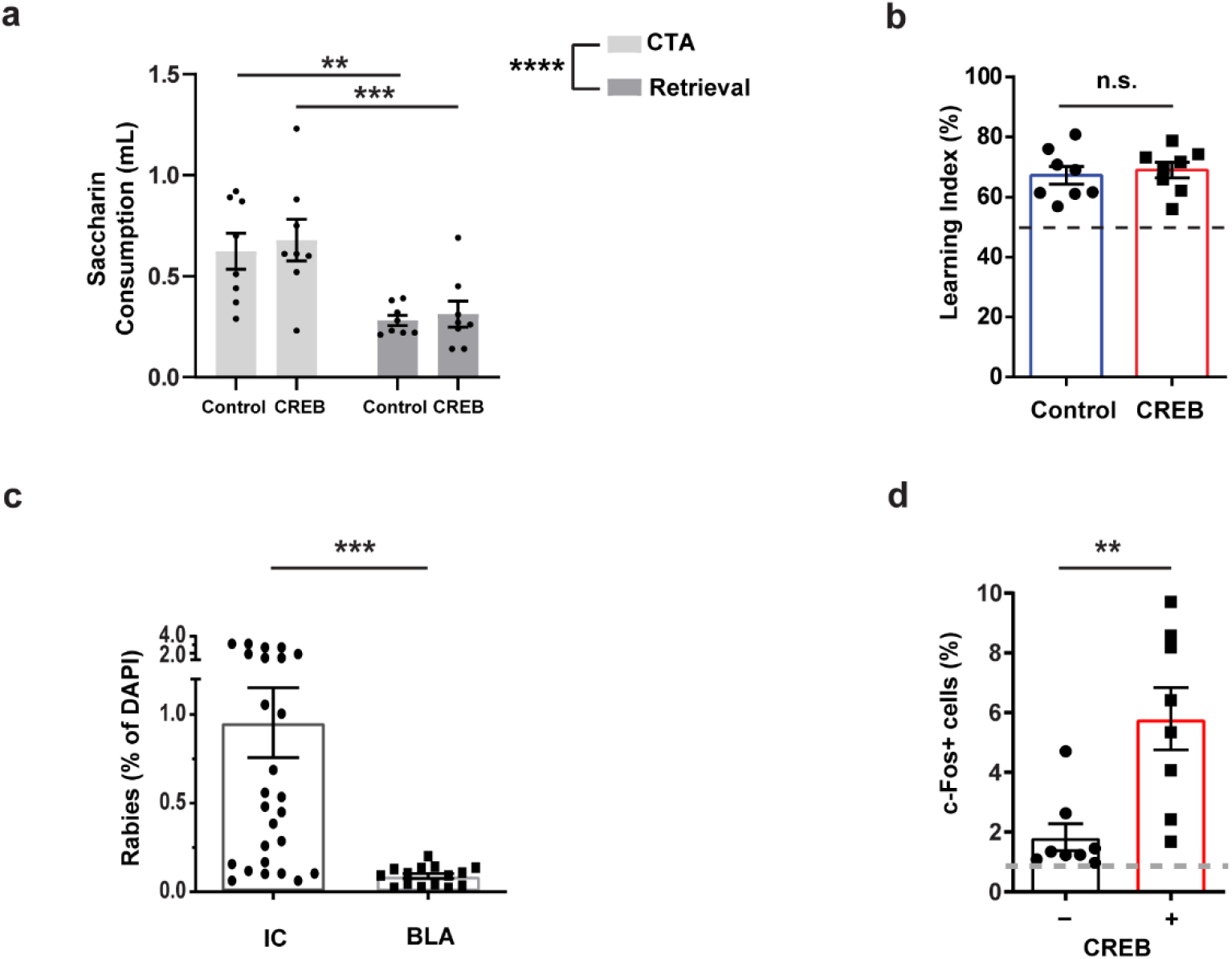
CTA with CREB expression in IC. **a**, Following CREB expression in IC, mice acquire CTA, reducing Saccharin consumption, in both CREB and control group (two-way RM ANOVA, F(1, 14) = 42.42, \*\*\*\**P*<0.0001, \*\*\**P*<0.001, **P<0.01). **b**, no difference in learning index between CREB and control mice. (unpaired t-test, P > 0.05). **c**, Percentage of rabies positive neurons from DAPI in IC and in BLA, where CRANE system was placed in BLA and IC, respectively (unpaired t-test, \*\*\**P*<0.001). **d**, Comparison of c-Fos expression in CREB-expressing neurons (CREB +; 6%) and neighboring neurons (CREB-; 1.42%) shows bias in memory allocation to vCREB neurons (unpaired t-test, \*\**P*<0.01). All results shown as mean ± s.e.m.

**Figure S4.**
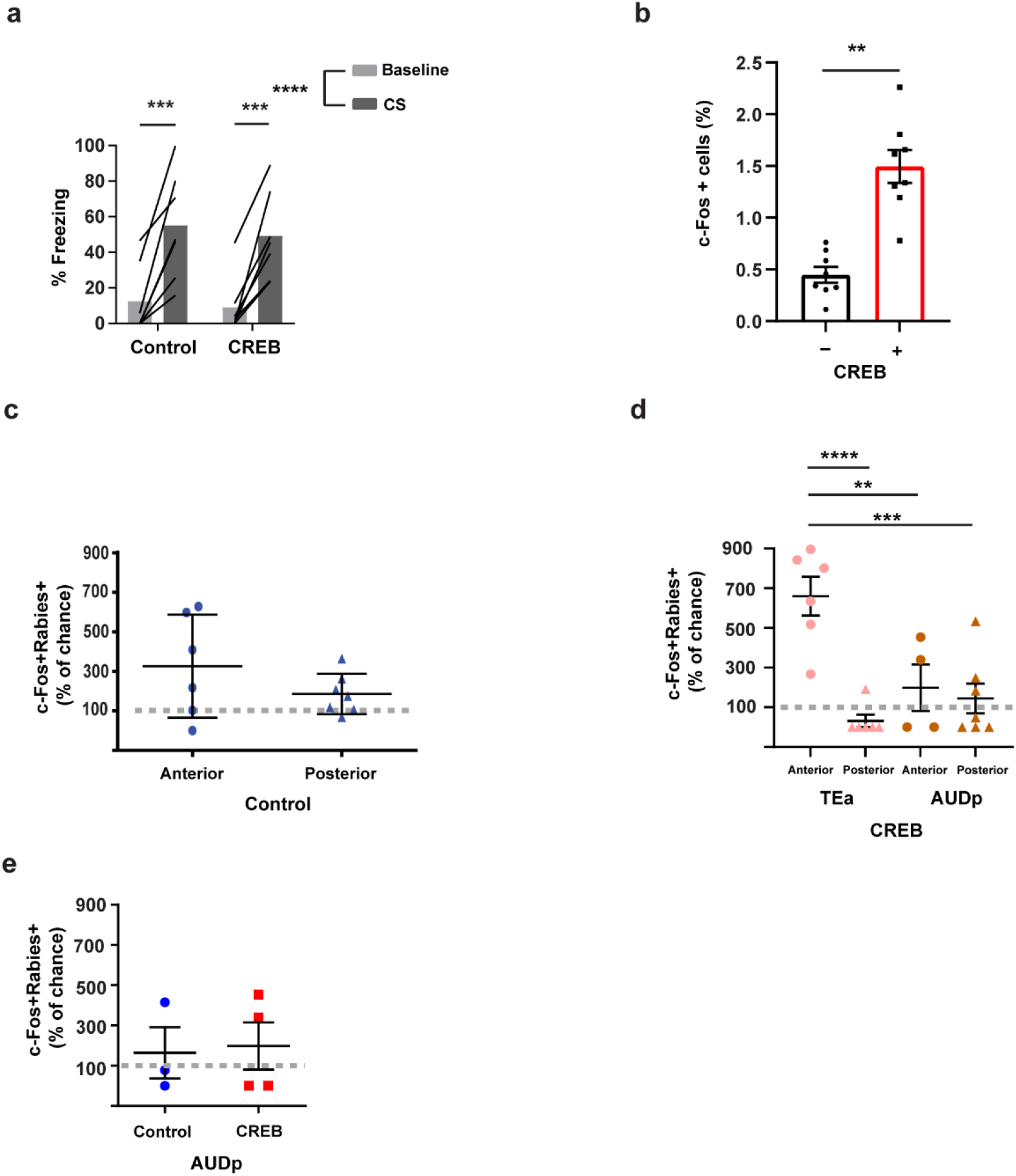
Characterization of CRANE and CREB expression in AFC. **a**, Comparing mice freezing levels between baseline and CS shows increased freezing in both CREB and Control (CFP) groups (two-way RM ANOVA, F (1, 12) = 60.93, \*\*\*\**P*<0.0001, \*\*\**P*<0.001). **b**, Comparison of c-Fos expression in CREB-expressing neurons (CREB +; 1.5%) and neighboring neurons (CREB -; 0.46%) shows bias in memory allocation to vCREB neurons (unpaired t-test, \*\**P*<0.01). **c,** No difference between anterior and posterior cortices in control group (unpaired t-test, *P*>0.05) or from chance levels (multiple one sample t-test, P>0.05). **d,** In the CREB group, the inter-regional network (i.e., Rabies positive) in the anterior temporal association area (‘TEa’) was highly activated (expressing c-Fos) in comparison to the posterior TEa and in comparison to the primary auditory cortex (‘AUDp’) (one-way ANOVA F(3,19)=12.23 *P*=0.0001; Dunnett’s multiple comparisons test \*\**P*<0.01, \*\*\**P*<0.001, \*\*\*\**P*<0.0001). **e**, No difference was found between CREB and Control groups, or from chance levels, in activation of the inter-regional network (c-Fos+Rabies+ neurons) in the primary auditory cortex (AUDp; Two-way ANOVA F(1,10)=0.0574 *P*=0.8155)

**Figure S5.**
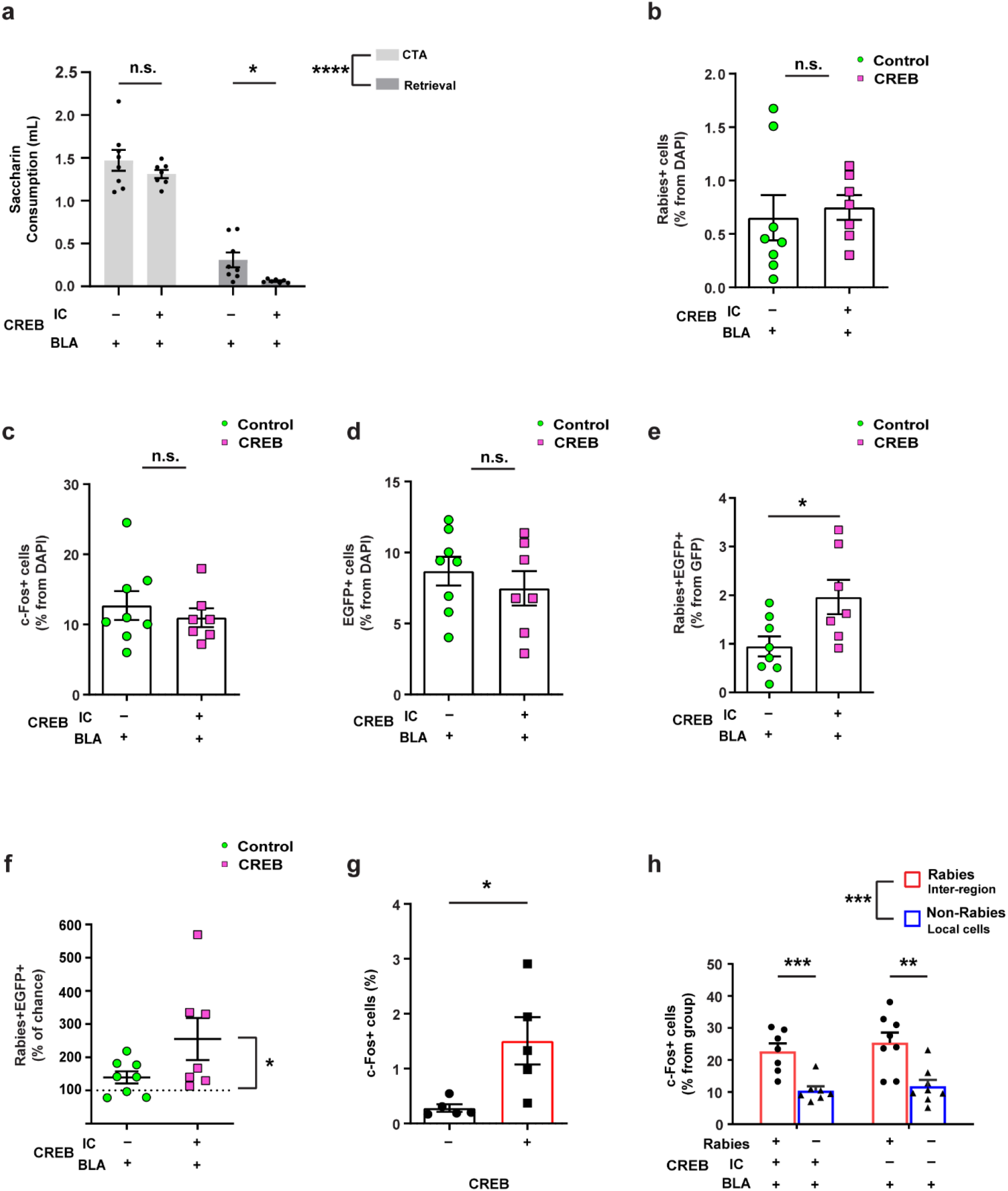
CREB expression in BLA and IC specifically enhances CTA memory retrieval. **a**, All groups expressed vCREB in BLA. CREB group also expressed vCREB in IC. Both groups acquire CTA with primary effect for CREB (no interaction; two-way RM ANOVA, F (1, 13) = 158.4 *P*<0.0001, F_CREB_(1,13) = 8.701 *P*=0.011, Šídák’s multiple comparisons test \*\*\*\**P*<0.0001). **b**-**d,** No difference between CREB and Control group was found in the expression of Rabies (**b**), c-Fos (**c**), or GFP (**d**). Unpaired t-test, *P*>0.05. **e,** Higher proportion of vCREB neurons in the IC connected to BLA vCREB neurons (Rabies+GFP+), in comparison to control group (GFP+; CFP in the IC; unpaired t-test, \**P*<0.05). **f,** vCREB neurons in the IC displayed higher than chance connectivity to BLA vCREB neurons (one-way ANOVA, F (2, 19) = 4.666, \**P*<0.05). **g,** Allocation of memory is biased to vCREB neurons in the BLA, in the CREB group (expressing vCREB both in the BLA and the IC). BLA CREB (CREB +; 1.5%) takes more c-Fos than neighbor cells (0.28%; unpaired t-test \**P*<0.05). **h**, Rabies neurons in the IC activated more than local cells in the IC (non-Rabies) in both groups (two-way ANOVA F(1,26)=29.38, \*\*\**P*<0.001; CREB \*\**P*<0.01; Control \*\*\**P*<0.001). All results shown as mean ± s.e.m.

